# RTFBSDB: an integrated framework for transcription factor binding site analysis

**DOI:** 10.1101/036053

**Authors:** Zhong Wang, Andre L. Martins, Charles G. Danko

## Abstract

Transcription factors (TFs) regulate complex programs of gene transcription by binding to short DNA sequence motifs. Here we introduce *rtfbsdb*, a unified framework that integrates a database of more than 65,000 TF binding motifs with tools to easily and efficiently scan target genome sequences. *Rtfbsdb* clusters motifs with similar DNA sequence specificities and optionally integrates RNA-seq or PRO-seq data to restrict analyses to motifs recognized by TFs expressed in the cell type of interest. Our package allows common analyses to be performed rapidly in an integrated environment.

**Availability:** *rtfbsdb* is available via github (https://github.com/Danko-Lab/rtfbsdb).

## 1. Introduction

Transcription factors (TFs) regulate complex programs of gene expression by modulating the rates of several steps early in transcription. TFs bind degenerate DNA sequence motifs, typically 3–20 bp, located in regulatory regions known as promoters, enhancers, and insulators. Identifying the coordinates of TF binding motifs within the genome is a crucial step in many genomic analyses. However, motif discovery is a challenging computational problem owing to the short lengths and high degeneracy of TF binding motifs.

Using experimentally derived sources of TF binding is one strategy to improve accuracy by constraining the motif discovery problem to known binding sequences. This strategy requires extensive knowledge about the DNA sequence specificities of TFs, which have historically been time-consuming to measure experimentally. Recently, high throughput experimental approaches have allowed the systematic discovery of motifs for thousands of TFs [1–3]. Moreover, strategies to impute binding motifs using TF amino-acid sequences extend these resources to most species with a sequenced genome [1]. These developments make the use of known TF binding motifs a powerful strategy in many common applications.

Here we introduce *rtfbsdb*, an open-source pipeline for transcription factor binding site (TFBS) identification and analysis, which integrates experimentally derived TF binding data for thousands of TFs. Unlike other TFBS identification tools, *rtfbsdb* integrates high-throughput measurements of gene expression for TFs associated with each motif. For downstream TFBS scanning and identification, *rtfbsdb* uses the *rtfbs* package [4], a highly flexible and efficient implementation of many TFBS scanning tasks. Many common and complex analyses can be solved by *rtfbsdb* in as little as a single line of R. We demonstrate *rtfbsdb* using genomic data from the ENCODE project.

## 2. Package Description

### Description of the rtfbsdb package

The *rtfbsdb* package is an open-source package for R which automates many common tasks in TFBS discovery and analysis. The flowchart for a typical analysis using *rtfbsdb* is presented in Fig. 1. To begin an analysis, users import a large database of experimentally defined TF binding motifs. We used the Catalog of Inferred Sequence Binding Preferences (Cis-BP) database, which integrates more than 65,000 motifs from >25 distinct experimental sources [1]. Cis-BP also includes imputed motifs for non-model organisms and for paralogs of well-characterized TFs. In addition to Cis-BP, *rtfbsdb* supports motifs from a variety of different sources and integrates data seamlessly in R.

**Fig. 1:**
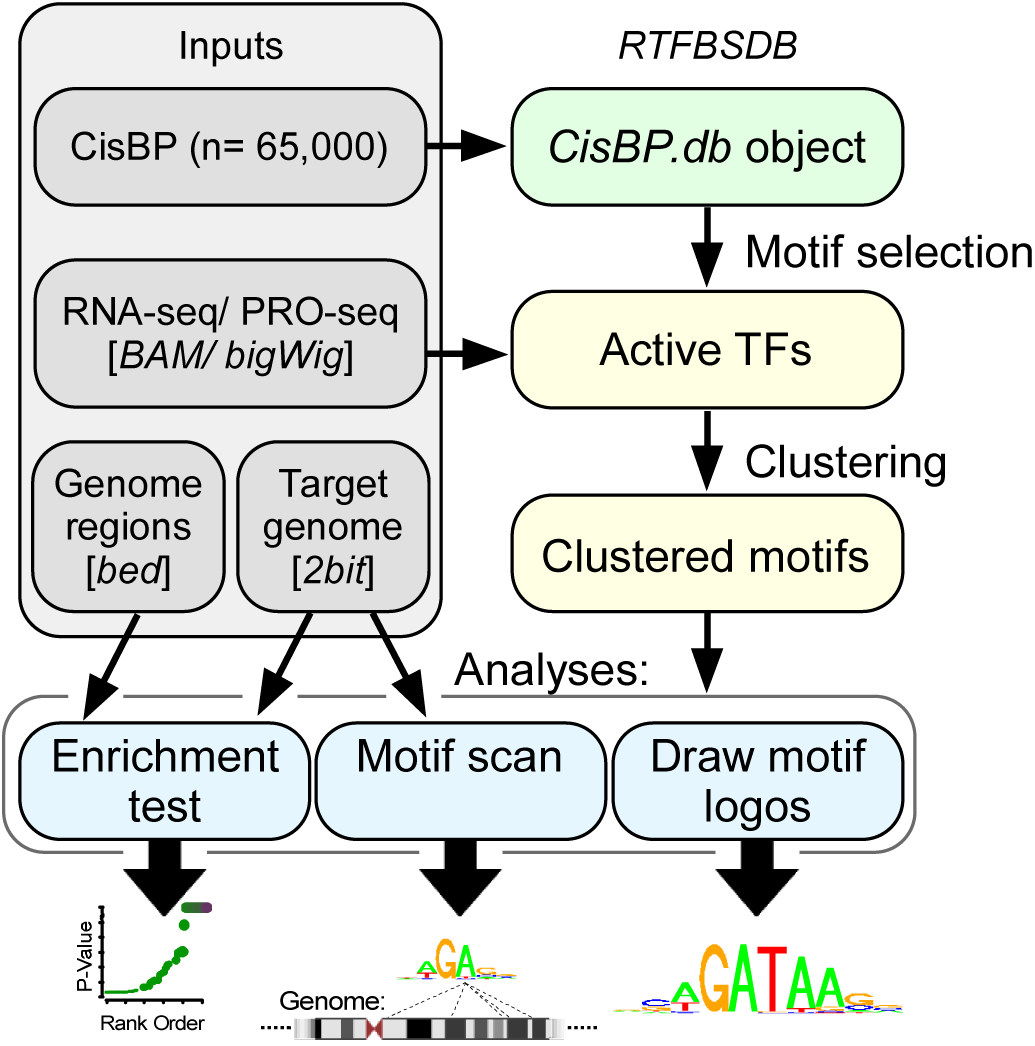
Schematic illustrating the *rtfbsdb* workflow. Motifs are loaded into a *CisBP.db* object in R using an automated web scraper that imports data directly from the Cis-BP database (green). The set of motifs is reduced to those most applicable for analysis (yellow, right side) by removing TFs that are not expressed in the cell system of interest, and subsequently grouping motifs recognizing similar DNA sequences by clustering. The final set of motifs can be used to complete several common tasks in genomics (blue, bottom row), including testing a set of DNA sequences for enriched motifs, scanning a target genome, or visualizing motifs as sequence logos.

DNA binding specificities are often highly similar between different TFs. Duplicate motifs reduce the interpretability of downstream analyses and can inappropriately decrease statistical power when using more stringent corrections for multiple hypothesis testing. We provide tools to focus analyses on motifs that are directly suitable for the user's application. Many analyses benefit from focusing on the subset of motifs for which the cognate TF is expressed in the cell type of interest. For this task, *rtfbsdb* integrates gene expression data collected using high-throughput sequencing approaches, including RNA-seq, PRO-seq, or related assays. *Rtfbsdb* estimates transcriptional activities of the TF associated with each motif in Cis-BP, which contains ENSEMBL IDs for each motif, and implements a statistical test [5] to identify those TFs which are expressed in the cell type or tissue of interest. To remove remaining redundant entries from *rtfbsdb* we cluster motifs based on their DNA sequence specificities using an affinity propagation clustering algorithm provided in the APCluster package [6]. After clustering, the similarities between each pair of motifs can be visualized as a heatmap (Fig. 2) and images of the motif logos within each cluster can be visualized in R or written to disk as a PDF file. The result of these pruning and clustering steps is to tailor the repertoire of motifs analyzed for the user's application.

**Fig. 2:**
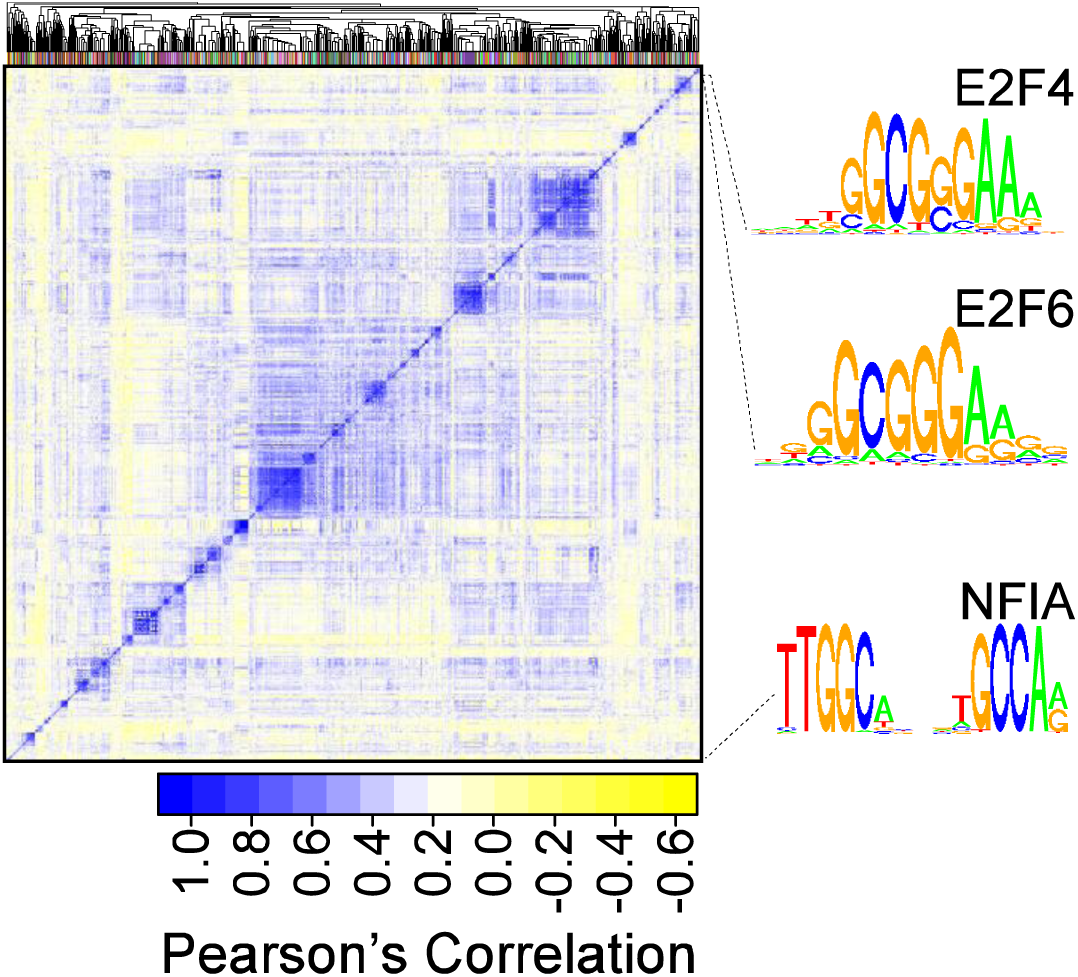
Correlation between motifs with similar DNA binding specificities. A correlation matrix grouping 1,964 motifs recognized by human TFs into 625 clusters using an affinity propagation clustering algorithm. The color scale represents Perason's correlation between the DNA sequence specificity represented by each motif.

After reading and filtering motifs, an *rtfbsdb* object can then be used to solve two classes of problem that are common in genomics. First, a common analysis is to identify the location of motif matches with a known DNA binding specificity across a target genome. Although this challenge is addressed by FIMO [7] and other applications, a notable advantage of *rtfbsdb* is that a database of experimentally derived motifs is integrated directly within the package. Moreover, our pipeline provides users with the option to write the coordinates of each motif directly to disk in a highly efficient compressed file format using bedops [8], enabling thousands of motifs to be scanned efficiently across large genomes.

Second, another common analysis task is to identify candidate TFs that putatively cause changes in gene expression. We provide tools to rapidly identify motifs enriched in test sequences compared to background. A systematic difference in dinucleotide composition between test and background sequences is the most common challenge with such an analysis. By default, *rtfbsdb* uses a resampling approach to identify background sequences with a similar distribution of GC content as test sequences. Additionally, *rtfbsdb* identifies motifs that are robustly enriched at several motif match score cutoff thresholds. Together, these innovations result in more reliable discriminative TFBS identification. To our knowledge, HOMER is the only other package that allows discriminative TFBS identification using experimentally derived TF binding motifs [9]. Compared to HOMER, *rtfbsdb* provides a larger repertoire of motifs, rigorously integrates TF expression levels using genomic data, and supports clustering motifs with similar DNA sequence specificities. Together, these advantages are likely to make *rtfbsdb* a more powerful and reliable tool for discriminative TFBS identification in many applications.

### Using multiple GC content groups decreases accuracy

A common step in TFBS identification is to divide loci into separate groups based on GC content and use a separate background model for each group. This strategy is assumed to accommodate systematic differences in GC content across the genome, and thus improve the specificity of motif matches. However, whether this practice results in superior TF binding site predictions has not been tested directly. We created an empirical test using publicly available data from the ENCODE project to determine whether dividing sequences into multiple GC content groups improves the accuracy of TFBS predictions. We used motif match scores to classify high-confidence DNase-I hypersensitives sites (DHS), defined as the intersection of DHS discovered using Duke and UW assays [10], as bound or unbound to its cognate TF. Chromatin immunopercipitation and sequencing (ChIP-seq) peaks from 21 TFs were used as a gold-standard set. Surprisingly, a background model constructed using all available sequences performs more accurately than dividing sequences into four separate groups by GC content in almost 90% of cases (Fig. 3). For example, identification of GATA2 and CTCF binding sites are 8.2% and 8.6% more accurate with only one GC content group. Thus, we conclude that using a single GC content group results in superior performance for the majority of TFs, and is the global default in *rtfbsdb*.

**Fig. 3:**
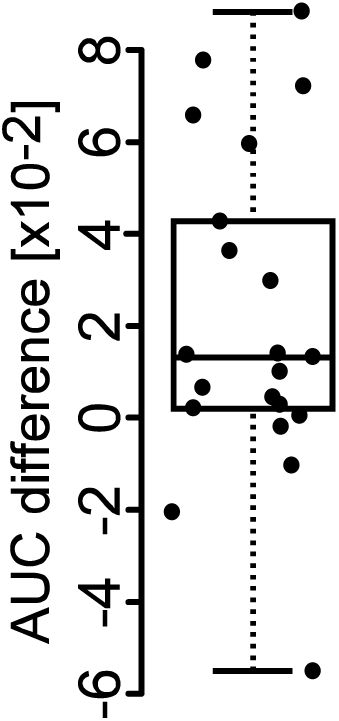
Accuracy of motif classification using 1 or 4 GC content groups. DNase-1 peaks in K562 cells were classified as ei-ther ‘bound’ or ‘unbound’ to 21 TFs using the motif match log-odds score. We com-puted the area under the receiver operat-ing characteristic curve (AUC) using ChIP-seq data in K562 cells as a gold-standard set, either with or without dividing se-quences into 4 separate GC content groups before scoring. The difference between AUCs when dividing sequences into either 1 or 4 separate GC content groups is indi-cated on the Y-axis. Individual points are shown, and the box-and-whiskers plot denotes the median, 25th and 75th per-centile, and maximum values.

## 3. Example

To demonstrate the utility of *rtfbsdb* we used motifs in Cis-BP to search ChIP-seq peaks discovered by ENCODE [11] for TFBS. We focused on ChIP-seq data profiling 97 TFs and co-factors in K562 cells. Forty-one of these are not represented by a motif in available databases, and these are largely comprised of either transcriptional co-repressors (e.g., HDAC1, EZH2, and KAP1) or general transcription factors (e.g., GTF3C2 and TAF1) without intrinsic sequence-specific DNA binding. Of the remaining 56 TFs the expected motif was recovered in 51 cases (91%), and was the most strikingly enriched motif in 42 (75%). For example, the motif corresponding to the transcriptional repressor REST was more than 50-fold enriched in ENCODE REST ChIP-seq peaks (Fig. 4). In the 25% of cases where the expected motif was not the most enriched, *rtfbsdb* typically recovered a motif corresponding to a known cofactor which likely recruits the expected TF to ChIP-seq peaks by proteinprotein interactions, in a process known as tethering. For example, peaks binding SP1 and SP2 were primarily enriched for NFYA and NFYB binding motifs, which represent a known TF tethering interaction [12]. Similarly, although EP300 contains a motif in CisBP, it is a transcriptional co-activator which is recruited to DNA by other TFs. We thus conclude that *rtfbsdb* returns the expected motif in real world test cases.

**Fig. 4:**
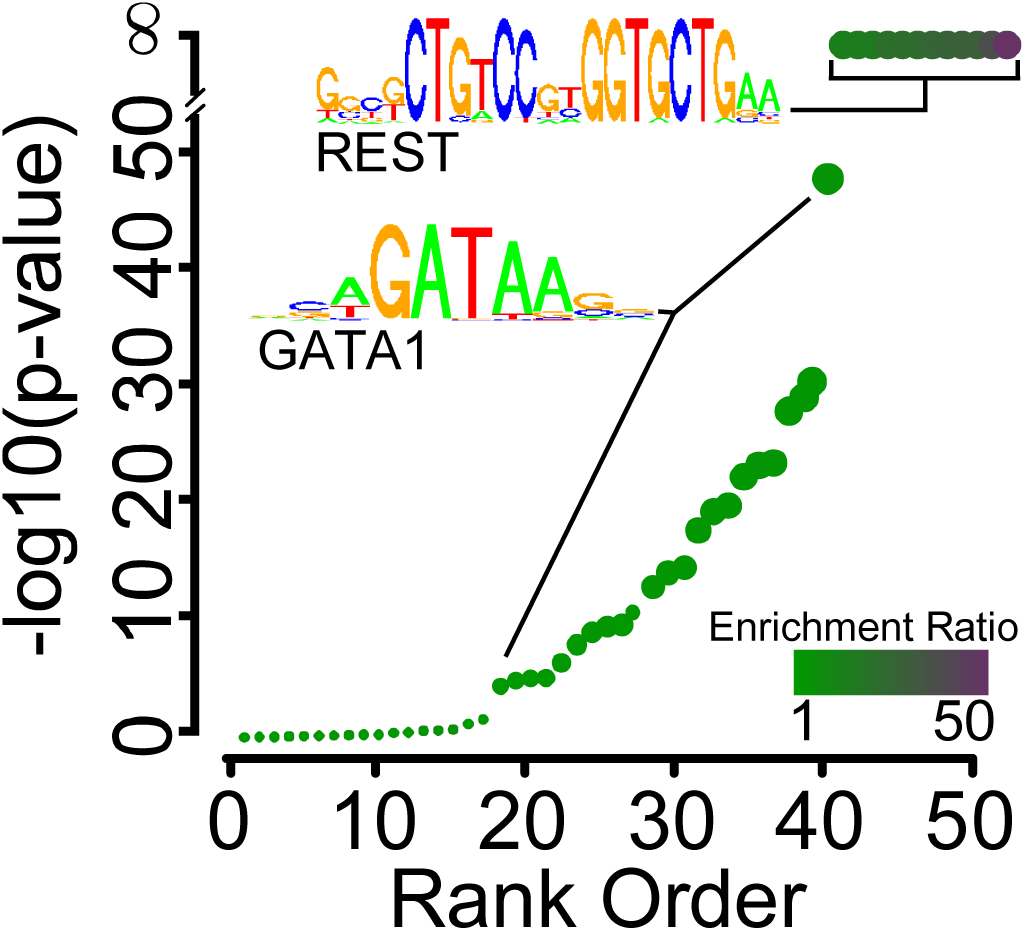
Enrichment of motifs in REST ChIP-seq peaks. The -log-10 p-value (Y-axis) as a function of the p-value rank order (X-axis) illustrates motifs enriched in ChIP-seq peaks binding the transcriptional repressor REST. The magnitude of enrichment is shown by the color scale and by the size of each point.

## 4. Acknowledgements

Work in this publication was supported by NHLBI (National Heart, Lung, and Blood Institute) UHL129958A to C.G.D. The content is solely the responsibility of the authors and does not necessarily represent the official views of the US National Institutes of Health.

